# Iron-Associated Mesenchymal Plasticity and Tumor Invasion in Glioblastoma

**DOI:** 10.64898/2025.12.22.695920

**Authors:** Christophe Petry, Ambra Stella Boecke, Aso Omer Mohammed, Nishitha Ghariwala, Vatsalkumar Jariwala, Mohammad Al Shhab, Oliver Buchholz, Ioannis Vasilikos, Julia M. Nakagawa, Andreas Vlachos, Jürgen Grauvogel, Mukesch J. Shah, Roland Rölz, Marco Prinz, Ulrich G. Hofmann, Jürgen Beck, Kevin Joseph, Vidhya M. Ravi

## Abstract

Glioblastoma (GBM) recurrence is driven by tumor cells that infiltrate surrounding brain tissue and evade surgical and therapeutic eradication. Although dysregulated iron handling is a recognized feature of the necrotic and hemorrhagic GBM tumor core, its relationship to invasive tumor cell states remains incompletely defined. Here, we investigated how iron-associated microenvironments relate to transcriptional programs of invasion in human GBM. Using multi-regional single-nucleus RNA sequencing of human GBM specimens encompassing tumor core, invasive front, and infiltrated cortex, we found that malignant cells from the tumor core exhibit coordinated upregulation of iron uptake and storage pathways together with invasion-associated gene programs. At the single-cell level, iron metabolism and invasion signatures were strongly correlated, defining a distinct core-enriched malignant subpopulation with mesenchymal-like transcriptional identity, stress-adaptive features, and angiogenic signaling. These iron-high/invasion-high cells aligned with mesenchymal-and astrocyte-like GBM states and were associated with unfavorable patient survival. Despite elevated oxidative stress and ferroptosis-associated transcriptional pressure, this population concurrently expressed anti-apoptotic and anti-ferroptotic regulators, consistent with an iron-tolerant invasive state.

To assess functional consequences of iron exposure, we modeled iron-rich conditions using non-cytotoxic particulate iron in patient-derived GBM cell lines and human organotypic cortical slice cultures. Iron exposure induced intracellular iron accumulation, oxidative stress responses, increased tumor cell motility in vitro, and enhanced invasion within intact human brain tissue. Collectively, these findings demonstrate that iron-rich tumor core niches are closely associated with mesenchymal plasticity and invasive behavior in GBM and support a role for iron-associated microenvironmental pressure in shaping invasive tumor cell states.

**Key Points:**

- Transcriptional profiling reveals that the GBM tumor core harbors malignant cells that strictly couple active iron metabolism with invasive programs.
- This iron-accumulating, Mesenchymal-like subpopulation is distinctively characterized by angiogenesis and stress resistance.
- Iron supplementation in vitro and human ex vivo models are sufficient to drive mesenchymal transition and significantly enhance tumor migration and tissue invasion.

**Importance of the Study:** Glioblastoma (GBM) recurrence is driven by highly invasive tumor cells that evade resection and resist therapy. Yet the microenvironmental pressures that push GBM cells into an invasive, therapy-resistant state remain poorly defined. Although iron dysregulation has been implicated across cancers, its role as a microenvironmental determinant of GBM invasion has never been demonstrated in physiologically relevant human systems. By integrating multi-regional single-nucleus RNA sequencing with functional validation in patient-derived GBM lines and human organotypic cortical slice cultures, we uncover a core-enriched, iron-associated mesenchymal program that co-segregates with invasion, stress adaptation, and angiogenic signaling. We further show that physiologically relevant iron exposure is sufficient to induce mesenchymal transition, enhance motility, and accelerate tissue invasion within human cortical architecture. These findings position iron as a selective driver that links the hemorrhagic, necrotic GBM core to the emergence of invasive subpopulations that seed recurrence. The data identify iron handling and stress-response pathways as actionable therapeutic vulnerabilities, providing a foundation for strategies that target metabolic resilience and iron-dependent invasive states in GBM.

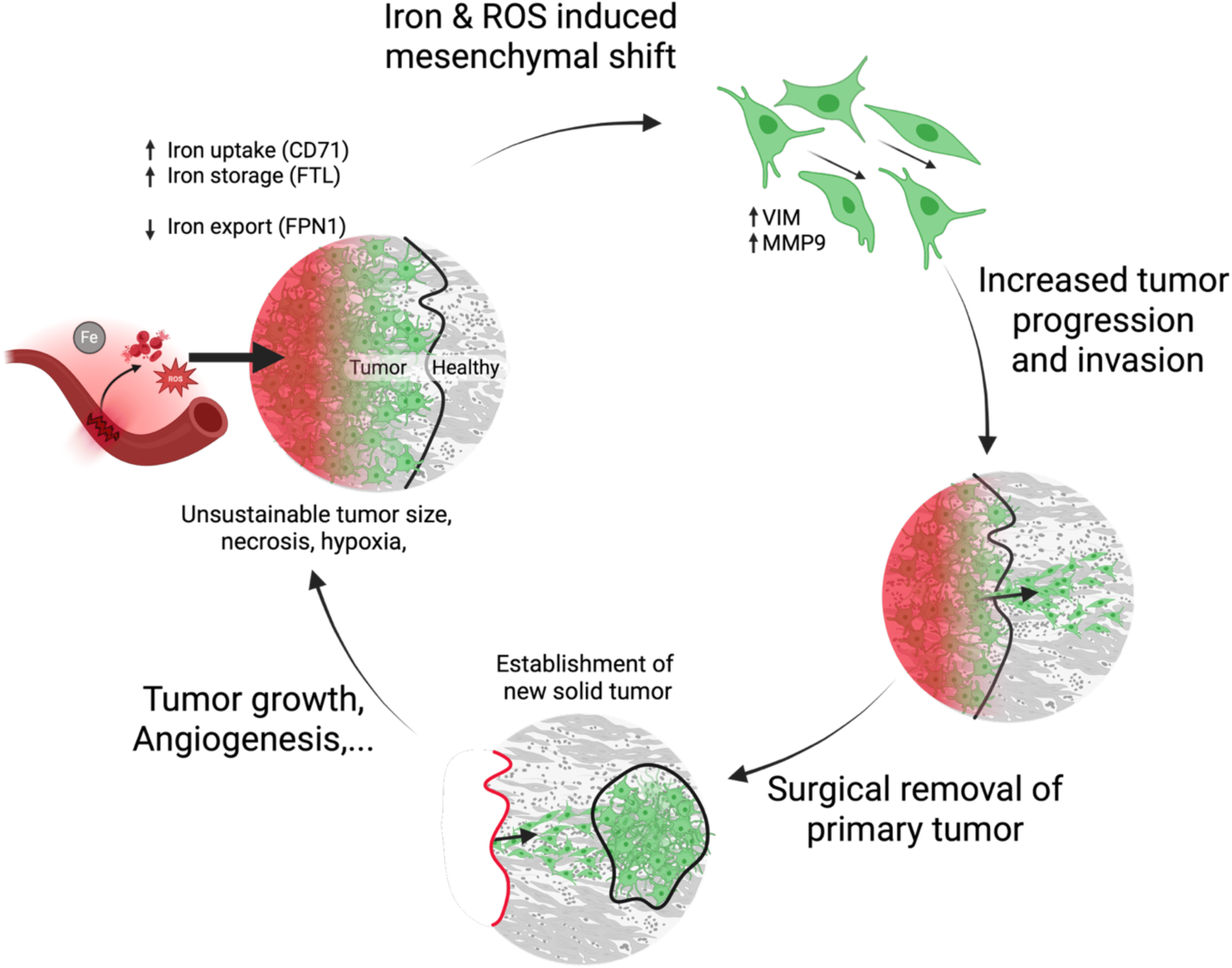

## Introduction

Glioblastoma (GBM) remains incurable because current treatments fail to eradicate the cells that have infiltrated tissue inevitably leading to recurrence ^1^. Despite extensive research efforts, the prognosis remains poor due to diffuse infiltration, treatment resistance, and tumor progression, often at the margin of resection^2,3^. GBM cells are highly heterogeneous, therefore understanding how some cells become invasive, and therapy resistant will help understand the cellular phenotype that escape the margin of resection enabling survival despite therapeutic intervention.

Iron is essential for fundamental cellular functions including DNA synthesis, energy metabolism, and oxidative phosphorylation^4^. However, excess labile iron is toxic, requiring controlled balancing of absorption, utilization, storage, and recycling known as iron homeostasis ^5^. Many cancers exploit this axis; mutations and transcriptional reprogramming in iron-regulating pathways enhance iron import and retention to fuel proliferation, metabolic rewiring, and survival under stress ^5,6^. This “iron addicted” phenotype, has been associated with increased proliferation ^7^, evasion of cell death^8^, and metabolic reprogramming in a range of tumor types ^9^. While the role that iron plays in GBM proliferation and therapy resistance have been explored, its direct contribution to invasion has not been systematically investigated.

In GBM, multiple features of the tumor microenvironment (TME) converge to create regions of high iron burden to which the tumor cells respond by modifying their iron handling and homeostasis ^10,11^. The GBM TME is shaped by regions of vascular leakage, microhemorrhages, hypoxia, and necrosis, resulting in sites of high iron concentration, and intense oxidative stress associated with chromosomal instability and mutations^12,13^. GBM stem-like cells (GSCs) that drive many malignant features are uniquely adapted to thrive in cyto-toxic micro-environments and exhibit a distinct upregulation of iron metabolism genes, enabling sustained proliferation under metabolic and oxidative stress ^11,14,15^. It is critical to understand the adaptive changes in the core that is thought to mediate the escape of tumor cells from a hostile environment, enabling them to infiltrate healthy cortex where they can revert to a more proliferative phenotype and seed recurrence ^16,17^

In this study, we investigate the link between iron metabolism and GBM tumor progression and invasion. We address this question by integrating single-nucleus RNA (snRNA) sequencing from human GBM core, invasive front, and infiltrated cortex with functional validation in patient-derived GBM cell lines and an ex vivo human cortical slice model. We identify a malignant subpopulation that concurrently upregulates iron metabolism and invasion signatures, enriched in the tumor core and transcriptionally aligned with mesenchymal-like states. We then demonstrate experimentally that exposure to physiologically relevant, non-toxic iron levels is sufficient to induce mesenchymal transition, enhance motility, and promote tissue invasion within human cortical microenvironments. Together, these findings reveal iron as an active microenvironmental driver of GBM mesenchymal plasticity and invasion, positioning the iron-rich tumor core as a metabolic niche that orchestrates the emergence and dissemination of aggressive, therapy-resistant tumor cells.

## Results

### Single-nucleus RNA sequencing links iron metabolism to mesenchymal, invasive programs in GBM

To define how iron metabolism varies across tumor regions, we generated an integrated snRNA-seq dataset of 49,000 nuclei from seven GBM patients, including contrast-enhancing tumor core, non-enhancing invasive front (**Fig. 1A**), and infiltrated cortex. After lineage-based annotation and copy-number profiling, 4,812 malignant nuclei were retained for downstream analysis (**Fig. 1B–C, Suppl. S1A-I**). As expected, we found the highest number of GBM cells in the tumor core, with declining numbers in invasive front and infiltrated cortex (**Suppl. Fig. S2J**).

**Figure 1:**
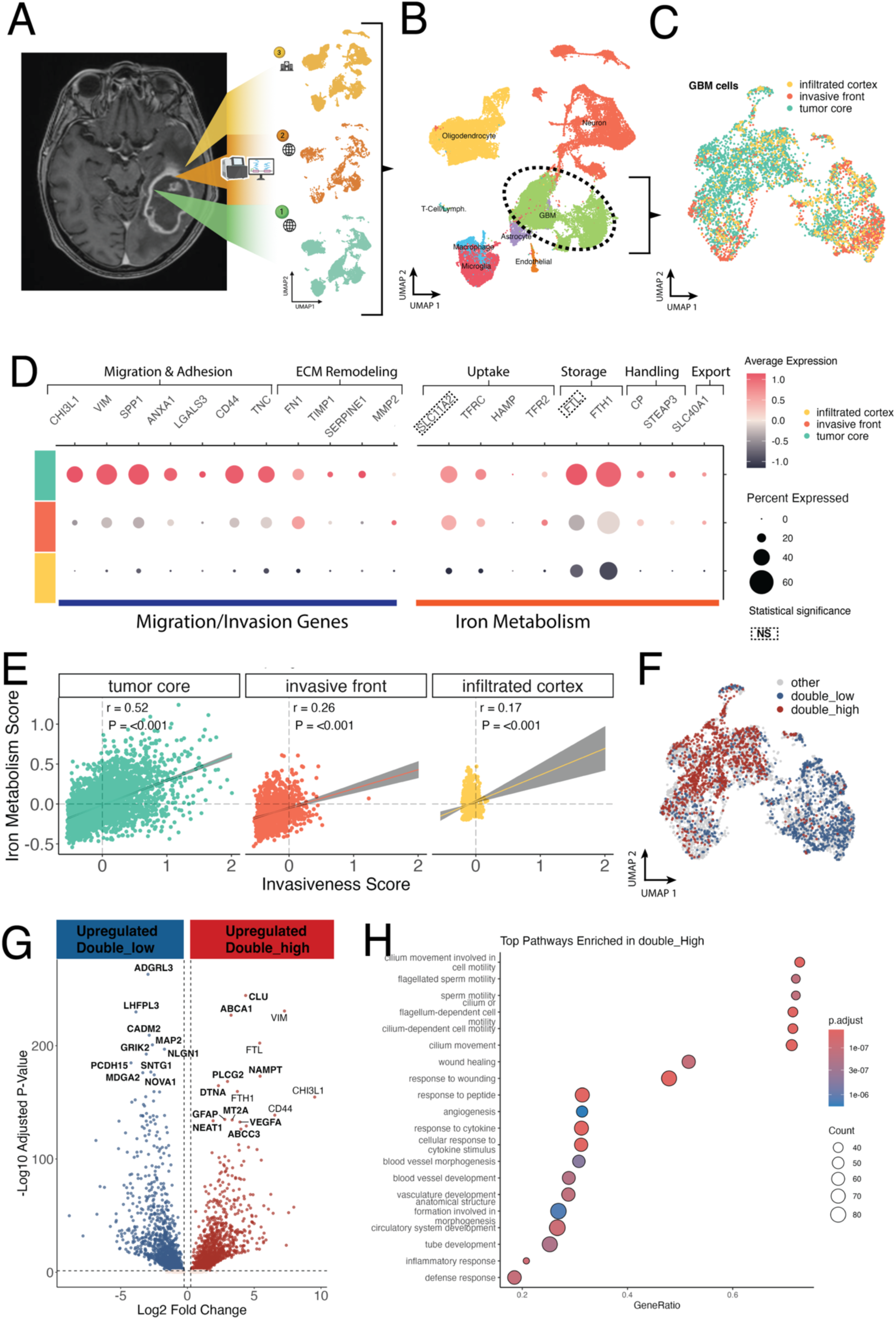
Iron metabolism and invasiveness are closely associated in the GBM tumor core. (**A**) Schematic showing single nucleus RNA sequencing samples originating from different regions of the tumor. Public data from tumor core (1. contrast enhancing region, N = 4) and invasive border (2. Non-enhancing region, N = 3) and in-house sequencing from the infiltrative cortex (3., N = 3). (**B**) Cell annotation of major brain cell types and identified GBM cells used for further analysis. (**C**) UMAP of the GBM cells colored based on the region of origin. (**D**) Dot plot showing the average expression level as well as percentage of cells expressing genes related to migration/invasion (left) and iron metabolism (right) depending on region of origin. Dashed boxes show non-significant genes; all other genes were significantly (p < 0.0001) upregulated in the core compared to invasive front and infiltrated cortex (**E**) Pearsońs correlation between iron metabolism score and invasion score (based on gene sets in D) depending on region of origin. Pearsońs r and p-value indicated in the plot (**F**) Cells were labelled based on their combined expression of genes related to iron metabolism and invasion. Cells that score “high” for both gene sets are termed double_high in contrast to double_low cells. Colored UMAP showing the location of double_high and double_low cells. (**G**) Volcano plot showing the differentially expressed genes between double_high and double_low cells. Negative values represent upregulated genes in double_low and positive values show upregulated genes in double_high cells. Fold change (0.25) and adjusted p-value (0.05) cutoffs are shown with dashed lines. 10 most highly upregulated genes per group are shown in bold (genes part of the defining gene set are in fine print). (**H**) Pathway analysis showing upregulated pathways that define the double_high population.

We first assessed whether the tumor core showed evidence of high iron use by scoring each cell for expression of general iron metabolism genes (uptake: *TFRC*, *SLC11A2*; storage: *FTH1*, *FTL*; reduction/handling: *STEAP3*, *CP* and export: *SLC40A1*) relevant in GBM ^23^. Core derived GBM cells expressed significantly higher levels of iron uptake and storage genes compared to cells from the invasive front and infiltrated cortex, accompanied by reduced iron export **(Fig. 1D, right).** We next assessed an invasion score defined by genes involved in cell adhesion and matrix remodeling (*CHI3L1*, *VIM*, *SPP1*, *ANXA1*, *LGAL*, *CD44*, *TNC*, *FN1*, *TIMP1*, *MMP2*, *SERPINE1*) ^24–26^. The same spatial pattern emerged: invasion associated transcripts were highest in cells from the core and diminished toward the outer boundaries of the infiltrated cortex (**Fig. 1D, left**). These findings characterize the tumor core as a micro-environment with active iron metabolism and invasion programs. This is consistent with our current understanding that the tumor core is highly populated with mesenchymal and stem-like GBM cells, known for their invasiveness and iron dependency, respectively ^11,27^.

To investigate whether this association also holds at the single-cell level, we scored each cell based on its expression of both iron metabolism and invasion genes. The scores were correlated within GBM cells, most strongly in the core (Pearson’s r = 0.52), then the invasive front (r = 0.28), and least in the infiltrated cortex (r = 0.16) (**Fig. 1E**), suggesting a potential cell-intrinsic link between both gene networks. To gain a better understanding of cells that concurrently express iron and invasion associated genes; to get a better understanding of the cells of interest we classified all cells based on their expression of iron and invasion genes into low/medium/high for each score (**Suppl. S2C**). Double^HIGH^ (iron^HIGH^/invasion^HIGH^) and double^LOW^ (iron^LOW^/invasion^LOW^) cells segregated on the UMAP, indicating transcriptionally distinct subpopulations (**Fig. 1F**). Indeed, double^HIGH^ cells were mainly associated with the MES- and AC-like state (**Suppl. Fig. S2D**), which are linked to distinct modes of GBM invasion ^28,29^. MES- and AC-like cells made up most subtypes in the core with OPC- and NPC-like cells dominating the invasive front and infiltrative cortex (**Suppl. Fig. S2G**). Differential gene expression analysis between double^HIGH^ and double^LOW^ cells showed that the former are defined by expression of genes associated with malignancy, stress/survival (*NEAT1*, *GFAP*, *PLCG2*, *DTNA*, *CLU*, *NAMPT*, *ABCA1*, *MT2A*) and angiogenesis (*VEGFA*) (**Fig. 1G, right**). Gene set enrichment analysis (GSEA) confirmed enrichment for motility, stress-response, and angiogenesis pathways (**Fig. 1H**). In contrast, double-low cells mainly occupied the NPC- and OPC-like states (**Suppl. S2D**), and were most represented in the invasive front, then core and absent in the infiltrative cortex (**Suppl. S2E**). These cells expressed genes for adhesion (*ADGRL3*, *NLGN1*, CADM2, PCDH15), neuronal structure (*MAP2*, *LHFPL3*, *SNTG1*), and signaling (*GRIK2*, *NOVA1*, *MDGA2*), alongside pathways consistent with neuronal features and cell cycle activity (**Suppl. S2F; Fig. 1G, left**).

We used the GBM validation cohort from “The Human Protein Atlas” ^30^ to check the influence of the differentially expressed genes on patient survival. The top upregulated genes in Double^HIGH^ cells were associated with significantly worse patient survival if highly expressed in the tumor (**e**xcept for *ABCA1*, *PLCG2*, *GFAP*) and the most highly differentially expressed gene *CLU*, also a prognostic marker for unfavorable patient survival. On the other hand, top upregulated genes in Double^LOW^ cells were associated with significantly better patient survival if highly expressed in the tumor (except for *LHFPL3*, *SNTG1*) and *ADGRL3*, *MDGA2* and *PCDH15* are potential prognostic marker for favorable patient survival. Underlining the difference in malignancy between these two groups.

We next assessed whether high iron accumulation is associated with apoptosis- and ferroptosis-related transcriptional programs. Cells originating from the tumor core and invasive front exhibited elevated expression of apoptotic genes compared with the infiltrated cortex. Notably, MES-like tumor cells concurrently displayed signatures linked to apoptosis and ferroptosis while also scoring highly for anti-apoptotic (*BCL2L1*, *BCL2*, *MCL1*) and anti-ferroptotic regulators (*SLC7A11*, *GPX4*) (**Suppl. Fig. S2H**).

Collectively, these data indicate that tumor core cells are exposed to iron-rich conditions and respond through coordinated upregulation of iron-handling pathways. Invasive transcriptional programs are tightly coupled to iron metabolism, both at the regional level and within individual tumor cells. Cells that co-express iron-associated and invasion-related genes represent a particularly aggressive population, characterized by mesenchymal identity, stress resistance, and pro-angiogenic features. This adaptive state likely enables mesenchymal tumor cells to exploit elevated intracellular iron while evading ferroptotic cell death, thereby positioning iron as a key regulator of GBM invasion originating from the tumor core.

### Acute particulate iron loading increases GBM motility and mesenchymal features in vitro

Having identified a transcriptionally distinct iron-accumulating, mesenchymal-like subpopulation enriched in the tumor core, we next sought to test whether iron availability directly alters tumor cell behavior. We modelled an iron-rich milieu using dextran-coated “nanoflower” maghemite IONPs due to their biocompatibility and established use in biomedical applications and it also reflects particulate iron released from erythrocytes during microhemorrhage (**Suppl. Fig S3A**) ^51^, We first titrated particle size (**50, 70, 100 nm**) and concentration (2.5, 5, 10 mg Fe/mL) for 24 h to identify a non-cytotoxic condition. Apoptosis increased at higher doses, especially with 100 nm particles at 5–10 mg Fe/mL (Suppl. Fig. S3B). All functional assays therefore used 70 nm at 2.5 mg Fe/mL (**Suppl. Fig. S3B**). To validate our model, we stained iron metabolism markers. At 2.5 mg Fe/mL (70 nm, 16 h), iron exposure significantly increased the expression of iron uptake and storage proteins like transferrin receptor (TFRC**/**CD71) and ferritin light chain (FTL), while iron export (ferroportin, FPN1) showed a statistically detectable but small decrease (**Fig. 2A**). Exposure to iron also caused a significant increase in HMOX1 indicating that the cells are responding to ROS (**Suppl. Fig. S3C**). This confirms effective iron loading and ROS production closely resembling the tumor core environment, under non-toxic conditions.

**Figure 2.**
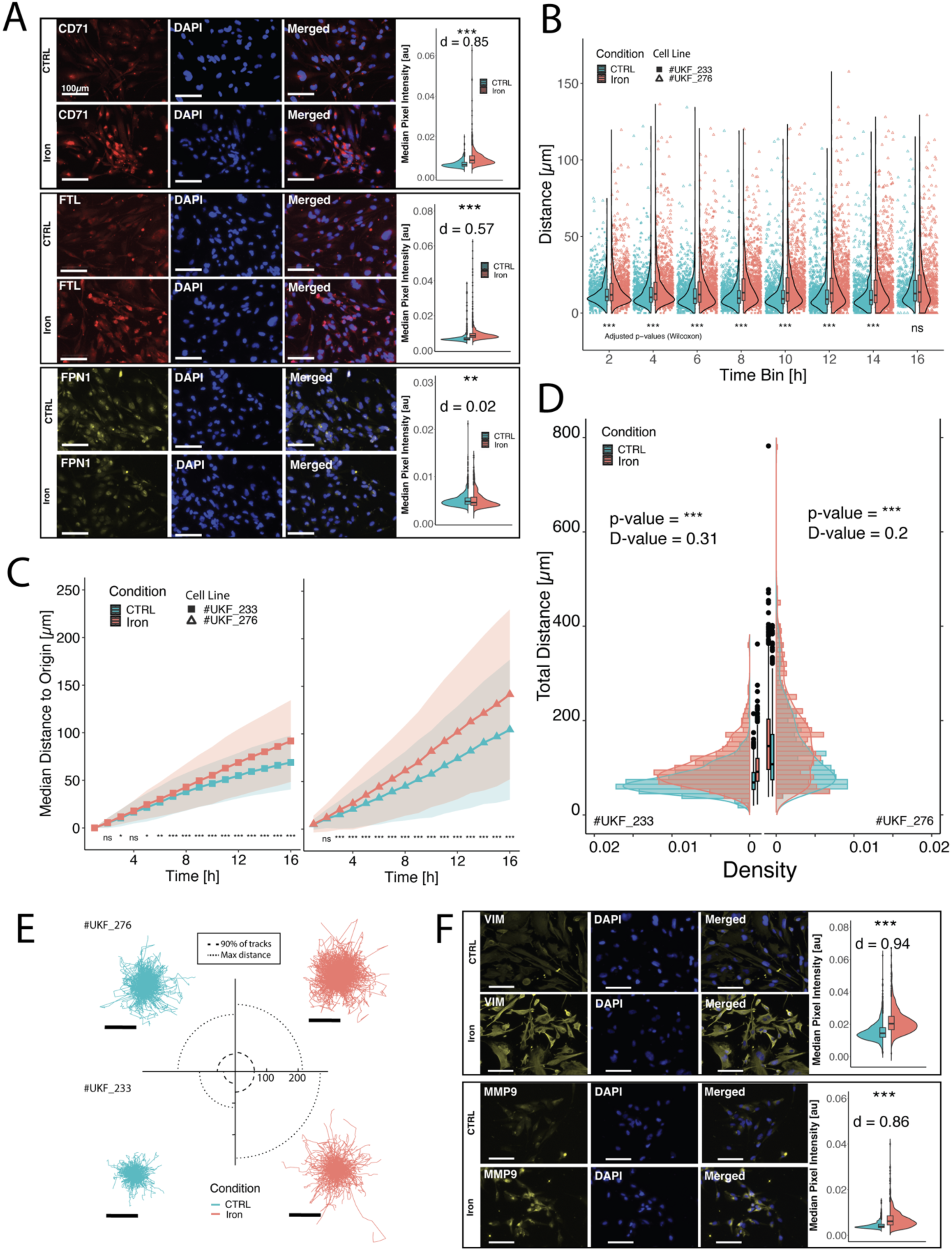
Iron exposure promotes a migratory phenotype in GBM cells in vitro. **(A)** Quantification of immunofluorescence stainings comparing median pixel intensity of cell bodies (violin plots) between conditions. Illustrative raw images of stainings for CD71 (transferring receptor protein 1; n_CTRL_ = 646, n_iron_ = 1029) (**top**), FTL (ferritin light chain; n_CTRL_ = 646, n_NP_ = 1029) (**middle**) and FPN1 (Ferroportin-1; n_CTRL_ = 968, n_iron_ = 762) (**bottom**). Scale bars are 100 µm and statistics (p-values and effect size d are shown above the violin plots. Distances migrated by GBM cells within 2-hour time bins over 16-hour exposure time, shown for both cell lines (#UKF_233 and #UKF_276) and treatment conditions (control and iron-exposed). Cumulative cell migration, representing the total distance the cells travelled over time. (**B-C**) Statistical evaluation by Wilcoxon rank sum test, adjusted for repeated measurement. (**D**) Distributions illustrating the total migration distances per cell across the full imaging period (16 h) in both conditions and cell lines. Additionally, boxplots summarize central tendencies and spread the data. Differences between distributions were assessed using the Kolmogorov–Smirnov test, with D-values indicating the degree of divergence. (**E**) Migration tracks showing the trajectories of individual cells over the 16-hour period relative to a common point of origin for each cell line and condition. In the center of the plot for comparison dashed circles indicate the radius encompassing 90% of all recorded cell positions; dotted circles represent the maximum observed displacement from the point of origin. Scale bars 100µm. (**F**) Quantification of median cell body pixel intensity (violin plots) and illustrative raw images of Vimentin (**top**) (VIM; n_CTRL_ = 1431, n_iron_ = 775) (**bottom**) and MMP9 (n_CTRL_ = 577, n_iron_ = 757). Scale bars are 100 µm and statistics (p-values and effect size d) are indicated above the plots. (**B-E**) Data pooled from 12 technical replicates for #UKF_233 (n = 959 cells; n_CTRL_ = 471, n_iron_ = 488) and 8 technical replicates for #UKF_276 (n = 1240 cells; control = 469, iron = 771).

Single-cell time-lapse imaging (16 h) in two patient-derived lines ***(#****UKF233**, #**UKF276*) demonstrated higher motility upon iron exposure (**Fig. 2B–C**). Temporally resolved (2h) analyses shows that this effect is fast with significant differences arising within 2h (**Fig. 2B**). Although there are line-specific differences in kinetics: #UKF276 exhibited an early increase (∼2 h) that attenuated after ∼12 h, whereas #UKF233 showed a delayed (∼6 h) but sustained rise (**Suppl. Fig. S3G**). The primary endpoint, cumulative path length per cell at 16 h, increased under iron in both lines. Total-distance distributions shifted towards greater distances with exposure to iron (Kolmogorov–Smirnov tests), with greater variance in #UKF276 (**Fig. 2D**). Cell trajectory plots from a common origin show hints of only a subpopulation of #UKF276 responding with increased migration and the bulk of the population remains relatively unaffected, whereas #UKF233 displayed a more uniform population-wide shift (**Fig. 2E**). Proteomic evidence points to a shift towards a mesenchymal phenotype, with expression of markers of ECM-remodeling markers (VIM, MMP9), being increased in the cells exposed to iron (**Fig. 2F**).

We aimed to model an iron rich environment using dextran-coated IONPs, the clinically used particulate form rather than soluble Fe salts; all in-vitro readouts therefore address particulate iron loading within a non-cytotoxic window. We show that supplementation with IONPs can model high iron environments shown by increased iron metabolism *in-vitro,* leading to active iron accumulation in cells and mild oxidative stress. This is accompanied by increased migration likely due to a shift towards a more mesenchymal state, which aligns with our in-silico findings. However, whether this translates to invasive behavior requires further experiments.

### Tumor progression and invasion is enhanced in *ex-vivo* human cortex model under conditions of elevated iron

To determine whether the in-vitro motility increase observed with particulate iron translates to human tissue we employed organotypic cortical slice cultures and assessed GBM growth and invasion after a single, non-cytotoxic iron exposure (70 nm, 2.5 mg Fe/mL) at inoculation. Given its more pronounced migratory response to iron in vitro, we selected #UKF_276 GBM cell line for these experiments to minimize the use of scarce human cortical tissue. Cortical slices from 3 tissue donors were inoculated with GBM cells and in the iron-rich condition, the injection solution was additionally supplemented with iron (2.5 mg Fe /mL) resulting in a one-time exposure at the time of injection (N = 3, n = 16). Tumor growth was monitored over 6 days of culturing. Tumor progression was tracked by fluorescent microscopy. Through imaging-based analysis, different metrics that reflect tumor growth, invasion and tissue colonization were extracted from the images (**Fig. 3A**).

**Figure 3:**
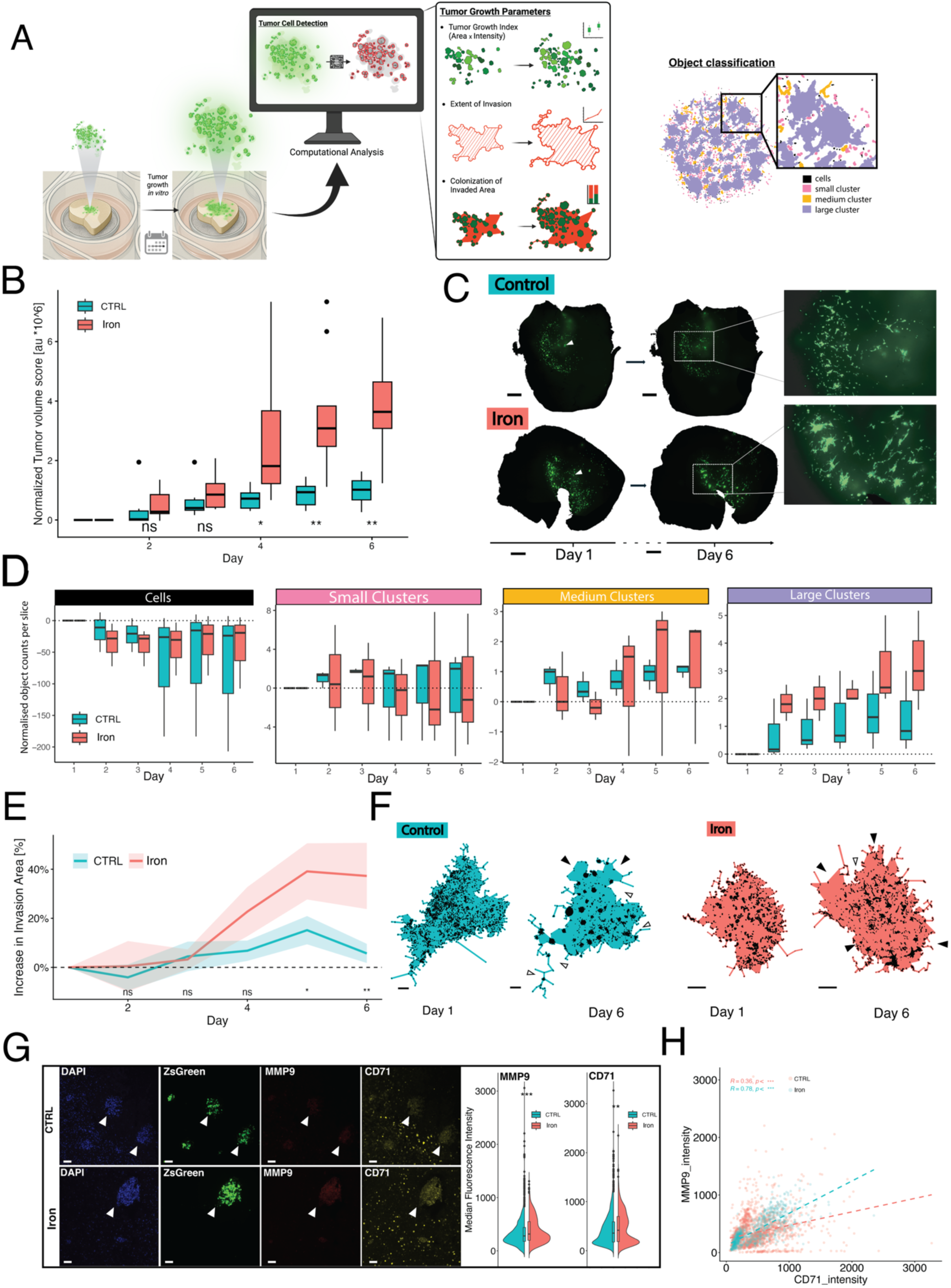
**Increased levels of iron in the cortical environment enhance tumor growth and invasion**. (**A**) Schematic representation of human cortical slice culture and extracted parameters of tumor progression. Left: Human cortical slices are inoculated with GBM cells and cultured over 6 days. Middle: Fluorescent tumor cells are detected, and their area and intensity are evaluated to compute the tumor volume score, extent of invasion and tissue colonization. Right: Illustrating the stratification of detected objects by coloring based on size. Shown are individual cells (black) small/medium/large clusters (pink, yellow and violet respectively). (**B**) Boxplots showing tumor volume score over 6 days of culturing normalized to day 1 (day of injection). Adjusted (FDR) p-values indicated below. (**C**) Two illustrative slices with tumor cells (slice masks in black and cells in green) showing increased cluster formation compared to control. Scale bar 1mm (**D**) Boxplots of object counts per slice normalized to day 1 showing the changes over days. (**E**) Median (line) and standard error of the mean (ribbon) of extent of invasion over time. Adjusted p-values indicated below. (**F**) Two examples illustrating the evolution of the tumor invasion area (colored based on condition) from day 1 to day 6. Invasive fronts are highlighted with a filled arrow, tumor shrinkage is highlighted with empty arrows, scale bars 500 µm. (**G**) Immunofluorescence images from slice culture (N = 3) injected with GBM cell (green) stained for MMP9 and CD71. Scale bar 100µm and white arrow heads highlight location of tumor clusters for ease of comparison (left). Quantification of mean pixel intensity of the tumor cells (right), (n_Control_ = 1962; n_Iron_ = 854). (**H**) Correlation between the cells fluorescence of MMP9 and CD71, comparing control vs iron exposed. Spearman correlation and p-value indicated above

First, we quantified tumor growth by computing a tumor volume score, a metric estimating 3D growth in the tissue (see *Slice culture proliferation analysis*). In accordance with our in vitro findings, we found that the tumor volume is significantly increased in the iron condition as early as 4 days post treatment (**Fig. 3B**). Tumor volume is highly varied in the iron condition, in some days reaching levels 6 times higher compared to the median of the control (**Figure 3B**). Visual inspection confirmed denser tumor structures and increased cluster formation in iron-treated slices (**Fig. 3C**). To better understand these patterns, we segregated detected tumor areas by size to distinguish isolated cells and different cluster sizes (**Fig. 3A right**). This allowed us to determine how the number of individual cells and the degree of cellular clustering changes over time. We could see that clustering is favored when GBM cells are exposed to excess iron since the number of large clusters per slice was significantly increased, indicating enhanced tumor cell aggregation and local proliferation (**Fig. 3C-D**).

Next, we sought to evaluate how well GBM cells invade the surrounding tissue. Beyond the increase of tumor volume, invasion has the additional component of cells penetrating healthy tissue. To track invasion, we determined the invasion area as the region bounded by a concave hull encompassing all detected tumor cells (**Fig. 3A**). While increased invasion was evident as early as day 4, a statistically significant difference emerged only by day 5 (**Fig. 3E**). Whereas tumor area plateaued under control conditions, slices exposed to iron continued to expand, showing a median increase of 37% in tumor area by day 6 (CTRL: 6%). In contrast, tumors in the control group often regressed, with visible cell loss and limited infiltration (**Fig. 3F**). Immunofluorescence stainings of slice culture tissue align with our cell culture findings in that iron exposed GBM cells increase their iron uptake (CD71) and MMP9 expression (**Fig. 3G**). Moreover, iron uptake and MMP9 strongly correlate in the iron treated slices compared to control (**Fig. 3H**).

In disease progression the step following invasion is tissue colonization, where cells switch from migration to proliferation and establish a new tumor core ^17^. To evaluate the ability of tumor cells to colonize newly invaded regions, we compared tumor volume to invasion area over time. This ratio provides an estimate of how effectively cells colonize tissue, switching from migratory to proliferative behavior after invading tissue. Although iron-treated slices consistently showed higher colonization efficiency, the differences were not statistically significant, likely due to high inter-sample variability (**Suppl. Fig. S4D-E**). In summary, even a single exposure to a physiologically relevant concentration of iron significantly accelerates GBM progression in an ex vivo human cortex model even days after. Iron promotes increased tumor expansion, tissue invasion, and local colonization, potentially through iron-mediated increases in MMP9 expression.

## Discussion

In this study, we identify iron availability as a characteristic feature of the glioblastoma (GBM) tumor core that is closely associated with mesenchymal plasticity and invasive behavior. By integrating multi-regional single-nucleus RNA sequencing with functional validation in patient-derived GBM cell lines and human organotypic cortical slice cultures, we show that iron-rich tumor regions harbor a distinct mesenchymal-like tumor cell population that couples active iron metabolism with invasion-associated transcriptional programs.

First, we investigated the role of iron in GBM invasion based on extensive evidence linking iron metabolism to malignant behavior and epithelial–mesenchymal transition (EMT)–associated invasion across multiple cancer types ^31,32^. Using multi-regional single-nucleus RNA sequencing from the tumor core, invasive front, and infiltrated cortex, we found that GBM cells in the tumor core are characterized by concurrent enrichment of iron metabolism and invasion-associated transcriptional programs (**Fig. 1A–D**). Active iron accumulation has been described in several malignancies and is thought to support tumor growth and progression ^33^. In GBM, this iron-associated transcriptional program progressively diminished with increasing distance from the tumor core, mirroring gradients of vascular abnormality within the tumor microenvironment and implicating hemorrhage, leaky vasculature, and iron-scavenging glioma stem-like cells (GSCs) as potential contributors to local iron enrichment ^34,11^. Importantly, beyond regional enrichment, iron metabolism and invasion programs were co-expressed at the single-cell level, indicating a cell-intrinsic coupling between these gene networks (**Fig. 1D, E**).

To further characterize this population, we focused on GBM cells that concurrently expressed high levels of iron metabolism and invasion-associated genes (Double^HIGH^) (**Suppl. Fig. S2E**). This transcriptional state corresponds to a highly aggressive phenotype enriched in the tumor core. Although apoptotic and ferroptosis-related programs were active in core and invasive-front regions, mesenchymal GBM cells exhibited pronounced upregulation of anti-apoptotic and anti-ferroptotic regulators, underscoring their capacity to withstand metabolic and oxidative stress (**Suppl. Fig. S2H**). This observation is consistent with prior reports demonstrating that the tumor core is dominated by mesenchymal- and astrocyte-like GBM states with enhanced stress tolerance and therapy resistance ^28, 35,36^ including reduced susceptibility to ferroptosis and a reliance on iron-dependent metabolic pathways ^37,38^. Such features likely reflect adaptive responses to iron-rich conditions, in which iron serves both as an essential cofactor for metabolic and enzymatic processes and as a source of oxidative stress that selectively favors resilient, invasion-competent tumor cells ^16^. In contrast, Double^LOW^ cells, defined by low expression of both iron metabolism and invasion programs, were predominantly NPC- and OPC-like and exhibited a progenitor-like, proliferative transcriptional profile characterized by cell–cell adhesion and network integration, consistent with reduced invasive capacity (**Fig. 1G**, **Suppl. Fig. S2F**). To functionally validate our in silico observations, we next examined the effects of elevated iron availability on GBM cell behavior using complementary in vitro and ex vivo human-based models. Monoculture assays enabled controlled assessment of cell-intrinsic responses to iron, whereas human organotypic cortical slice cultures allowed evaluation of tumor growth and invasion within a physiologically relevant tissue environment. We modeled an iron-rich microenvironment using dextran-coated iron oxide nanoparticles (IONPs; 70 nm, 2.5 mg Fe/mL), selected based on titration experiments to achieve non-cytotoxic intracellular iron loading (**Suppl. Fig. S3B**). This particulate iron model provides a controlled approximation of iron accumulation that may occur in regions of hemorrhage or necrosis while avoiding acute toxicity associated with soluble iron salts. Exposure to IONPs induced increased expression of iron uptake and storage markers, indicating intracellular iron accumulation (**Fig. 2A**), and phenocopied iron-associated transcriptional features observed in tumor core cells by snRNA-seq (**Fig. 1D**). Similar alterations in iron metabolism resulting in excess intracellular iron have been reported across multiple cancer types and are associated with enhanced metabolic capacity under conditions of oxidative stress ^39^. While increased iron availability supports metabolic and enzymatic demands, it also imposes persistent oxidative stress, which has been linked to genomic instability, tumor progression, and malignancy ^12,13^.

Functionally, iron accumulation enhanced multiple aspects of malignant behavior in GBM cells. Iron exposure increased mesenchymal and extracellular matrix remodeling markers and promoted tumor cell motility in vitro (**Fig. 2B–F**), consistent with prior studies showing that iron supplementation can induce mesenchymal transition and migration in diverse cancer contexts ^40,41^. Although these effects may be directly mediated by iron-dependent metabolic pathways, increased oxidative stress likely contributes as a parallel or interacting mechanism. Supporting this interpretation, iron exposure induced expression of the oxidative stress response gene HMOX1 (**Suppl. Fig. S3C**), a pattern also observed in tumor core transcriptomic data. In GBM, reactive oxygen species (ROS) signaling is known to promote migration and invasion through HIF-dependent mesenchymal programs ^42^, and iron-rich conditions may further amplify this effect, as HIF-1 activation can enhance iron uptake ^43^. Consistent with the in vitro findings, GBM cells inoculated into human cortical slice cultures and exposed to iron exhibited accelerated tumor progression, increased cluster formation, and enhanced tissue invasion over time (**Fig. 3B–F**). validating the relevance of iron-associated invasive programs in a human brain microenvironment. In iron-supplemented slices, tumor cells showed increased expression of iron uptake (CD71) and the invasion-associated protease MMP9, with a strong correlation between both markers (**Fig. 3G–H**). This association suggests a functional link between iron accumulation and invasive machinery, potentially mediated by redox-sensitive transcriptional regulators such as NF-κB and HIF-1α, both of which are known to regulate MMP9 expression ^44,45^.

The distinct response kinetics observed between monoculture and tissue-based models likely reflect the cellular complexity of cortical tissue, where non-tumor cells contribute to iron handling. Microglia and astrocytes are primary regulators of iron uptake, storage, and release in the brain ^46^. Depending on activation state, microglia can sequester iron or release it into the extracellular environment, particularly in anti-inflammatory (M2-like) states characterized by increased iron export and heme degradation^47,48^. Notably, regions enriched for mesenchymal-like tumor cells in GBM often contain abundant tumor-associated macrophages/microglia, and our single-nucleus data indicate that Double^HIGH^ tumor cells express genes implicated in microglial recruitment and polarization, including ABCA1 and CHI3L1 ^49^. These M2-like cells preferentially localize to perivascular and necrotic regions, potentially reinforcing localized iron-rich niches. Previous work using human organotypic slice cultures has similarly shown that GBM growth and invasion preferentially occur near reactive glial populations, reflecting the spatial organization of human tumors ^27,50^.

Together, these findings support a model in which iron acts as an important microenvironmental factor within the GBM tumor microenvironment that, alongside hypoxia and necrosis, contributes to selective enrichment of invasive and stress-resistant tumor cell states. Cells capable of adapting to iron-rich conditions acquire enhanced migratory and invasive potential, facilitating escape from the hostile tumor core. As tumor cells infiltrate surrounding brain tissue and environmental constraints are relieved, they may regain proliferative capacity, enabling re-establishment of tumor mass and perpetuation of disease progression.

In summary, these observations highlight iron handling as a relevant component of GBM aggressiveness and suggest that therapeutic or diagnostic strategies involving iron modulation should be carefully evaluated in the context of tumor invasion.

## Funding

This work was supported by BMBF (DiaQNOS: 1010 0665 01) and DFG MA 10605/1-1.

## Conflict of Interest

All authors declare no potential conflict of interest.

## Author Contributions

C.P.: Human Brain Slice Culture experiments; cell culture experiments; data analysis; sn-RNAseq analysis; Imaging; figure generation; writing/reviewing manuscript. A.B.: sn-RNAseq preprocessing (iCNV),. A.O.M.: primary cell line generations, sn-RNAseq library processing, illumina sequencing and data acquisition. N.G.: Immunofluorescence staining. V.J.: Immunofluorescence quantification. M.A.S: primary cell line generation, Tissue Collection, sn-RNAseq library processing. O.B.: cell culture experiments. I.V., A.V., J.G., M.J.S., J.N., R.R.: Neurosurgical brain tissue samples, review manuscript. M.P.: Neuropathology, review manuscript. U.G.H.: study supervision, review manuscript, grant acquisition for the study. J.B.: Study supervision, review manuscript, grant acquisition for the study. K.J: sn-RNAseq data curation and preprocessing, code verification, writing & reviewing manuscript, study supervision. V.M.R., study supervision, guidance of experiments, writing/revising manuscript, grant acquisition for the study. All authors were involved in reviewing the manuscript,

## Data availability

Public data is available at: “https://www.synapse.org/NEvCE_snRNAseq_PatelTessema”. In-house sequencing data will be shared on GEO (in progress). All custom scrips are available on GitHub repository: “https://github.com/ChrisPetry97/Iron-rich-tumor-core-niches-fuel-mesenchymal-invasion-in-glioblastoma”

## Acknowledgments

We thank Depro Das, Christian Munkel and Jonathan Goeldner for insightful discussions and technical guidance. We acknowledge BioRender.com for assistance with figure generation. We are grateful to all patients who generously donated tissue for this study. All R code used for data analysis was written, reviewed, and validated by the authors. Large language models (ChatGPT,OpenAI) were used in a very limited and supportive manner to assist with refactoring, and as well as to provide suggestions to improve clarity and grammar of the manuscript text. LLM was not used for data interpretation or to generate scientific conclusions. LLM was additionally used to generate a schematic illustration of a cortical slice in culture included as part of Figure 3A.

## Materials and Methods

### Patient samples

The data presented in this study includes human data from publicly available snRNA sequencing data ^18^ from contrast enhancing and non-enhancing regions of the tumor combined with our own in-house snRNA sequencing of human infiltrated accessory cortex (#UKF541: male, 81yo temporal; #UKF613: male, 63yo, temporal; #UKF698: male, 61yo, temporal) excised to access deeper GBM pathologies. Human GBM and cortical access tissue were obtained from patients undergoing neurosurgical resection at the Medical Center – University of Freiburg after written informed consent, in accordance with the Declaration of Helsinki and institutional guidelines. The study was approved by the Ethics Committee of the University of Freiburg (*EK-Freiburg: 24-1100-S1*), which cover the use of infiltrated cortex for single-nucleus RNA sequencing and ex vivo slice cultures experiments.

### Sn-RNA sequencing prepratation and analysis

Frozen human cortical tissue was homogenized, lysed, purified and FACs filtered following the recommendations from 10x Genomics to yield high quality nuclei. Nuclei were then processed using the 10x Genomics Chromium Next GEM 3′ v3.1 workflow. Libraries were amplified and sequenced on an Illumina NextSeq 2000. Reads were aligned using the Cell Ranger pipeline (v7.1.0 and v8.0.0; 10x Genomics) with the GRCh38-2020-A or GRCh38-2024-A reference transcriptome (supplementary materials *Single nucleus sample preparation* and *sequencing*). Computational analysis was performed using R package Seurat (v5.3.0). Standard preprocessing steps were taken on each individual sample including normalizing, batch-correcting (SCTransform), regressing out mitochondrial RNA content and finally integration, dimensionality reduction and clustering was performed on the combined data set (supplementary materials *Single nucleus sequencing analysis-preprocessing*).

The first step of the downstream analysis is cell type annotation and the goal here is to identify the malignant cells. Cells were annotated based on canonical CNS markers and cells that did not clearly fit any particular cell type were potentially malignant and further checked for chromosomal abnormalities. 5838 cells were identified as GBM cells and reintegrated (supplementary materials *Single nucleus sequencing analysis – annotation*). We checked the expression of curated gene sets related to iron metabolism and tumor invasion, compiled from published literature. Cells were further labelled according to their expression level of iron and invasion genes (cells with high expression of both gene sets: Double^HIGH^; and cells with low expression of both Double^LOW^) and according to their GBM subtype using the gene sets from Neftel et al. 2019. Differential gene expression between Double^HIGH^ and Double^LOW^ populations was performed using Wilcoxon rank-sum tests. Genes were filtered with thresholds of log2 fold change > 0.25 and detection in at least 10% of cells. Pathway analysis was performed to check gene pathways that define Double^HIGH^ or Double^LOW^ populations (supplementary materials *Single nucleus sequencing analysis – GBM cells* and *iron metabolism and invasion*).

### Iron Oxide Nanoparticles

To emulate the gradual release of iron from iron-rich erythrocytes, we used Synomag®-D dextran coated iron oxide nanoparticles (IONPs) with a core size of 50 nm, 70 nm and 100 nm were tested. Before use IONPs were sterilized using a syringe filter with pore size 0.45 µm and heated to 37 °C (supplementary materials *Iron Oxide Nanoparticles*). After testing their cytotoxicity (supplementary materials *TUNEL assay*) 70nm at 2.5mg Fe/mL was chosen for in vitro and ex vivo experiments. For cell culture experiments, cells were exposed to a cell culture medium supplemented with IONPs at a concentration of 2.5 mg Fe/mL for 16h. For tissue culture experiments, cell solutions used for inoculation were supplemented with 2.5 mg Fe/mL.

### Cell tracking

Fluorescently labelled cells (#UKF233, #UKF276) as previously reported were plated on laminin-coated 96-well plates and left to settle overnight. Immediately before the experiment, cells received fresh medium (control or iron supplemented) and were imaged every 30min using an EVOS M7000 microscope with an onstage incubator for 16h at 10x magnification. Cells were tracked using the Fiji ^19^ plugin TrackMate7 ^20^ (supplementary materials *Cell culture, fluorescent microscopy* and *Cell tracking*).

### Human Brain Slice Cultures

Human cortical tissue directly from the operation room was prepared for long term culturing as previously described ^21,22^. In brief, human cortical tissue removed to access deeper pathologies was collected immediately from the operation room and processed into 300 µm thick slices and cultured for 6 days (supplementary materials *Human organotypic slice culture*). After one day slices are inoculated with 1 µL of a 10‘000 cells/µL cell solution containing fluorescently labelled cells (#UKF276) (and in the iron condition 2.5 mg Fe/mL) (supplementary materials *Slice culture injections*). Slices are imaged daily to track tumor growth. Main progression parameters include tumor volume score, which considers area and fluorescent intensity of tumor cells to estimate volume, tumor cluster formation, which tracks the evolution of cells and clusters over the days, tumor invasion, which follows the concave hull around all detected tumor cells, and finally tumor colonization, which monitors how much of the invaded area is populated by tumor cells. Supplementary materials *Slice culture tumor progression analysis / tumor cell and cluster analysis / invasion analysis / tissue colonization*.

### Immunofluorescence staining

Cells were treated as in *Cell tracking* except in 24 well plates with laminin coated glass coverslips and PFA fixed for 10 minutes at room temperature. OCT embedded PFA fixed cortical slices were sectioned to 20 µm and mounted on glass slides. Staining cells and sections followed the same basic protocol of blocking, permeabilizing, primary and secondary antibodies with variations on incubation times and permeabilization (Supplementary materials *Immunofluorescence stainings*). Comparative probes were imaged with constant imaging parameters and median pixel intensity (of soma or nucleus depending on antibody target) was compared between conditions (Supplementary materials *Immunofluorescence quantification*).

## Statistical analysis

Statistical analysis was performed using R. Statistical comparison of medians was done using the non-parametric Wilcoxon rank sum test. P-values for repeated measurements were adjusted using FDR. Counts (like different detected objects in slice culture experiments) were compared using chi-squared test. P-values are indicated as follows: p-value<0.05 as *; p-value<0.01 as **; p-value<0.001 as ***). Effect size (Coheńs D) is computed for large sample sizes. Correlations were evaluated using Spearmańs correlation.

## Notes

https://www.synapse.org/NEvCE_snRNAseq_PatelTessema

### Competing Interest Statement

The authors have declared no competing interest.

