## Supplemental Info for "Iron-Associated Mesenchymal Plasticity and Tumor Invasion in Glioblastoma"

### Supplementary methods and figures

#### Single nucleus sample preparation

To prepare high-quality single nuclei for sequencing, snap frozen tissue samples were thawed on ice, and any residual liquid was removed. For fresh frozen samples embedded in OCT, a wash with nuclease-free water was performed to remove excess embedding material. Tissue was homogenized using Nuclei EZ Lysis Buffer, followed by incubation on ice and centrifugation at 500g for 5 minutes at 4°C to isolate nuclei. The now pelleted nuclei were resuspended in fresh lysis buffer, incubated on ice, and centrifuged again to ensure thorough removal of cellular debris. After the final lysis step, the nuclei were resuspended in Nuclei Suspension Buffer with DAPI (DNSB) and filtered through a Flowmi Cell Strainer (70 µm) to remove large debris and aggregates. To further purify the sample, a sucrose gradient was prepared by layering the nuclei suspension mixed with 30% sucrose onto a sucrose base layer. This gradient was centrifuged at 13,000g for 30–45 minutes at 4 ℃, separating the nuclei from contaminants such as myelin or cellular debris. The nuclei pellet was carefully collected, resuspended in DNSB, and filtered through a 40-µm Flowmi Cell Strainer for additional debris removal. The purified nuclei were then sorted using fluorescence-activated cell sorting (FACS), where DAPI-positive nuclei were isolated. Following sorting, if needed, the nuclei were centrifuged, resuspended in the remaining buffer, and prepared for Chromium loading following the 10x Genomics protocol.

#### Single nuclei sequencing

Single-nucleus RNA sequencing (snRNA-seq) libraries were prepared using the Chromium Next GEM Single Cell 3’ Reagent Kits v3.1 (10x Genomics) according to the manufacturer’s protocol, optimized for fresh-frozen tissue. In brief, nuclei were isolated from acute fresh-frozen human cortex samples (as described above) and loaded into the Chromium Controller for single-nucleus separation and barcoding. Reverse transcription and cDNA amplification were performed with 12 PCR cycles. Sequencing libraries were constructed using 13 cycles of PCR amplification. Final libraries were sequenced on an Illumina NextSeq 2000 platform, yielding between ~85 and ~152 million reads per sample. Sequencing achieved high base quality (Q30 > 93% across barcodes, UMIs, and RNA reads), with a mean of 7,400–17,500 reads per cell and sequencing saturation ranging from 34.1% to 59.7%. Between 6,400 and 11,400 cells were captured per sample, with median detection of 698–1,087 genes and 897–1,563 UMIs per nucleus. Read alignment and transcript quantification were performed using the Cell Ranger pipeline (v7.1.0 and v8.0.0; 10x Genomics) with the GRCh38-2020-A or GRCh38-2024-A reference transcriptome. All datasets were processed using the "--include-introns" option to accommodate nuclear transcripts, enabling the inclusion of both exonic and intronic reads. Mapping rates to the genome exceeded 95%, with 53–61% of reads mapping confidently to the transcriptome. Antisense mapping rates ranged from 19.7% to 22.2%, consistent with expected performance in nuclear RNA-seq.

#### Single nuclei sequencing analysis-preprocessing

Single-nucleus RNA sequencing data was processed and analyzed using a custom Seurat (v4.0-v5.3) pipeline in R. Preprocessed objects (tumor and non-tumor cells) from a recent publication [89] were merged and then split by patient identity in preparation for our data to be included. To account for distant infiltration, we added in-house data from infiltrated cortex of three patients (#UKF541, #UKF613, and #UKF698). Each dataset was log-normalized individually, and the proportion of mitochondrial reads per cell was calculated. Data was then variance-stabilized using SCTransform for downstream analysis, and mitochondrial read percentage were regressed out. To prepare for integration, anchors were identified across samples using the first 30 principal components. The object was then integrated, and the integrated assay was scaled and subjected to dimensionality reduction (PCA and UMAP). Nearest neighbors of cells were computed (FindNeighbors()) and clustering was performed with a resolution of 0.5. Clustering was visualized using UMAP plots grouped by patient and region to verify proper integration and clustering. Data from Patel et al. (2024) is available online (https://www.synapse.org/NEvCE_snRNAseq_PatelTessema). Our in-house data will be uploaded to GEO.

#### Single nuclei sequencing analysis - annotation

After preprocessing our dataset includes a total of 48636 cells from tissue resected during GBM operations of 7 different patients. We labelled the cells based on the region of origin relative to the tumor. These regions include the tumor core (1. contrast enhancing region; n = 16830 cells), infiltrative front (2. non-enhancing region; n = 3629 cells) and our in-house sequenced infiltrated cortex (3.; n = 28177 cells). The main goal of the annotation step is to find the GBM cells which will be used for downstream analysis. To determine which cells in our SnRNA-seq data are GBM we checked for canonical cell marker genes (**Suppl. S1A-B**). Well defined clusters that show clear cell type markers were annotated (oligodendrocytes, microglia/macrophages, neurons and endothelial cells) and we termed suspected malignant cells as "unclassified" (Seurat clusters: 1, 2, 5-11, 16-22) (**Suppl. S1C**). Running CNV on those cells we found classic alterations usually found in GBM, of gain in chromosome 7 and loss in chromosome 10 (**Suppl. S1D, E**). Some of the initially conservatively annotated "unclassified" cells turned out to be neurons (clusters: 5, 6, 7, 9, 10, 18, 22), astrocytes (cluster: 16) and T-cells (cluster: 21) (**Fig. 1B**). Based on thresholding of CNV values relating to chromosome 7 and 10 we labelled cells as GBM (**Suppl. S1F**). Upon another close inspection of the CNVs of the remaining unclassified cells (clusters: 1, 2, 8, 11, 17) we found OPC cells which made up a cluster very low in CNV scores (**Suppl. S1G**). Finally, after removing the OPCs and reintegrating the GBM cells, these cells (n = 5838 cells) were subject to the following downstream analysis (**Fig. 1C**).

#### Single nucleus sequencing analysis – GBM cells

To prepare the newly subset GBM cells for further analysis they require some preprocessing. The RNA assay was set as default and datasets were split by patient, and samples with very few cells (<20; #UKF541 and Gr4-GBM-4) were excluded for appropriate integration. For each sample, genes with zero counts across all cells were removed prior to normalization to avoid issues during integration. Normalization and integration were performed as indicated above. In short, "SCTransform" was used for variance-stabilization and normalization. Integration features were selected across samples, datasets were prepared for integration with "PrepSCTIntegration" and anchors were identified using the first 20 principal components. Data was then integrated with a reduced k.weight parameter to account for variation in cell number, followed by dimensionality reduction (PCA, UMAP) and clustering with a shared nearest neighbor graph (resolution = 0.4). To classify GBM cells into transcriptional subtypes, we applied the gene signature framework described by Neftel et al. [61]. Subtype-specific signatures (MES1, MES2, AC, OPC, NPC1, NPC2) were scored per cell using AddModuleScore, and simplified into four major GBM programs (MES, AC, OPC, NPC). Each cell was assigned a subtype label according to the highest signature score. Subtype distributions were visualized on UMAP embeddings and by patient identity.

#### Single nucleus sequencing analysis – iron metabolism and invasion

Curated gene sets related to iron metabolism and tumor invasion were compiled from published literature. Iron metabolism-related genes (FTH1, FTL, TFRC, SLC40A1, STEAP3, SLC11A2, CP) were chosen to include iron uptake, storage, handling and export. Invasion-related gene sets were derived from both general cancer invasion markers (MMP2, MMP9, VIM, CDH2, FN1, STAT3, SRF) and glioblastoma (GBM)-specific invasion markers curated from prior studies (e.g., CHI3L1, CD44, SERPINE1, TNC, ANXA1, SPP1). We assess expression patterns of our defined gene sets "DotPlot()" and "FeaturePlot()" functions to examine regional differences and which GBM subtypes express our genes of interest (**Suppl Fig. S3A-B**). Genes with consistently low expression across samples (MMP9, RELB, BCL3, IL1B) were excluded from downstream analyses. Cells were scored based on their expression of iron (Iron metabolism score) and invasion genes (Invasion Score) from our data sets. Correlation between both scores was assessed using Spearman's rank correlation within each tumor region. Linear regression models were fit to visualize the relationship, and significance was evaluated with two-tailed correlation tests. Using our defined scores cells were classified into low, medium, and high expression based on tertile thresholds (**Suppl. Fig 3C**). Cells simultaneously high in both categories were classified as double_high, while those low in both were designated as double_low (the rest was labelled "other"). The labels were visualized on the UMAP to display distributions across cellular population.

To evaluate whether $Double^{HIGH}$ or $Double^{LOW}$ cells were enriched in specific regions, or subtypes, the relative proportion of each category was calculated per metadata slot. Stacked bar plots normalized to 100% were generated to visualize cell distribution across categories.

Differential expression between $Double^{HIGH}$ and $Double^{LOW}$ populations was performed using Wilcoxon rank-sum tests. Genes were filtered with thresholds of log2 fold change > 0.25 and detection in at least 10% of cells. Genes defining the iron and invasion modules which were used to define both groups were excluded. Significant differentially expressed genes (adjusted $p < 0.05$) were visualized via volcano plots and heatmaps. Top 10 genes in each group were labeled on the plot. Pathway analysis was performed on all genes to check gene pathways that define double_high or double_low populations.

#### Cell Culture

Patient derived glioblastoma (GBM) cells (#UKF233 and #UKF276) were genetically modified to express ZsGreen. Cell culture was carried out as previously described [49,83]. In brief, cells were cultured in adherent cell culture flasks at 37 °C and 5% CO2 in Minimum Essential Medium (MEM, GibcoTM, Thermo Fischer, Catalog number: 11090081) supplemented with 10% Fetal-calf serum (FCS) (P30-3306; PANBiotech GmbH, Aidenbach, Germany) and 1% Penicillin/Streptomycin (15140122, Thermo Fisher Scientific Inc., Waltham, MA, USA). Culture viability was ensured by frequently checking their confluence and the cells were passaged when necessary. All cells were used at a low passage number (<10).

#### Iron Oxide Nanoparticles

Synomag®-D dextran coated iron oxide nanoparticles (IONPs) with a core size of 50 nm (Micromod Partikeltechnologie GmbH, Catalog number 104-00-501), 70 nm (Catalog number

104-00-701) and 100 nm (Catalog number 104-00-102) were tested. The particles consist of a core of maghemite with „nanoflower" structure in a matrix of dextran. Before use IONPs were sterilized using a syringe filter with pore size 0.45 µm (Rotilab®-syringe filters, Catalog number P667.1) and heated to 37 °C. For cell culture experiments, cells were exposed to a cell culture medium supplemented with IONPs at a concentration of 2.5 mg Fe/mL for 16h. For tissue culture experiments cell solutions used for inoculation were supplemented with 2.5 mg Fe/mL.

#### TUNEL assay

A 96-multiwell plates was coated with laminin to improve cell adhesion. Next, the cells (#UKF233) were seeded at a density of 2500 cells per well in 200 µL of MEM to achieve the desired confluence. After 24 hours, the cells were adherent and ready for exposure to different concentrations (2.5, 5, 10 mg Fe / mL) and sizes (50, 70 and 100nm) of IONPs, triplicates were produced for each condition. Then all wells received fresh medium (control or with IONPs) and were incubated for 16h. After that an apoptosis assay (Terminal deoxynucleotidyl transferase-mediated dUTP-biotin nick end labelling [TUNEL]) assay (Cell Meter™ Live Cell TUNEL Apoptosis Assay Kit, AAT Bioquest, Catalog number 22844) was performed following the protocol in the instruction manual. In short, all components were brought to room temperature medium in the wells was removed, the working solution was added, and the plate was incubated for 60 minutes. The solution was removed, the wells washed twice with DPBS, and reaction buffer was added. After that we measured the fluorescence using a microplate reader (excitation 550 nm, emission 650 nm). Data about the apoptosis-related fluorescence in the two conditions were extracted. Experiments were carried out in triplicates.

#### Fluorescence Microscopy

For time-lapse microscopy, 96-multiwell plates were coated with laminin to improve cell adhesion. Next, the cells were seeded at a density of 2500 cells per well in 200 µL of MEM to achieve the desired confluence for imaging. After 24 hours, the cells were adherent and ready for imaging. Immediately before the start of imaging, all wells received fresh medium (control or with IONPs). Imaging was performed using the EVOS M7000 microscope (Thermo Fisher, Catalog number: AMF7000) with the EVOS Onstage Incubator option (Thermo Fisher, Catalog number: AMC1000), enabling automated long-term fluorescent imaging of multiple wells. Images were captured using a 10x objective over a 16-hour period. The imaging frequency was set to twice per hour, with an exposure time of 250 ms.

#### Cell tracking

The frames captured during live cell recording were combined into stacks and preprocessed using Fiji [84]. Images were converted to 8-bit, contrast was increased by 0.3% and the images were thresholded to binarize the cells. Leftover noise from the background and debris was filtered out using the median filter with a radius of 5 pixels. Cell movements over the frames were tracked using the ImageJ plugin TrackMate7 [85]. To detect individual cells the "Mask detector" was used as the preprocessing leaves only cell bodies to have non-zero pixel values. For tracking the "Linear Assignment Problem" (LAP) tracker [86] was used as it can handle track splitting (due to cell division). The frame-to-frame linking was set to link detected spots that are no more than 100 pixels apart, to avoid linking neighboring cells. Gap closing was disabled, and gap splitting was enabled for distances below 20 pixels. Finally, tracks were included in further analysis if they include detected spots from more than 90% of the recorded frames. This makes sure only reliably detected and tracked cells are included for further analysis. We verified the accuracy of our automated tracking by comparing it to manual tracking (**Suppl. Fig 3E**).

#### Immunofluorescence stainings

Glass coverslips (12 mm diameter) were pre-coated with laminin at 37 °C for 1 hour prior to use in a 24-well plate to improve cell adhesion. 50'000 #UKF_276 cells were seeded per well and cultured as stated in **Cell Culture** section. Triplicates were produced per condition for each antibody panel. On the following day, cells were treated with 2.5 mg Fe/mL IONPs for 16 hours. After treatment, cells were gently washed once with phosphate-buffered saline (PBS) and fixed with 4% formaldehyde (ROTI®Histofix, Carl Roth GmbH & Co. KG, order number 3105.1) in PBS for 10 minutes at room temperature. Following fixation, cells were gently washed three times with PBS for 5 minutes each. Coverslips were stored in the well plates covered in PBS and stored at 4°C overnight. For immunofluorescence staining, PBS was removed and replaced with blocking buffer solution (5% BSA and 0.1% Triton in PBS) for 1h at room temperature for permeabilization and to prevent unspecific antibody binding. Primary antibodies were diluted in blocking buffer at a concentration of 1:500 and applied to the cells for 2 hours at room temperature. After incubation, cells were washed three times with PBS for 10 minutes each on plate shaker on the lowest setting. Secondary antibodies (1:500), diluted in blocking buffer and containing DAPI (1:1000), were applied for 2 hours at room temperature in the dark. Cells were then washed three times with PBS for 10 minutes each. Following staining, coverslips were shortly dried and immediately mounted onto glass microscope slides using VECTASHIELD® Antifade Mounting Medium (©Vector Laboratories, SKU: H-1000-10). Slides were allowed to dry flat in the dark and stored at 4 °C until imaging.

For staining of cortical sections, frozen blocks were resectioned using cryostat at a thickness of 20 µm and stained following the same protocol except that incubation times were increased to 2 days for primary antibodies and 2h for secondary antibodies. Additionally, double the concentration (0.2% Triton) was used during permeabilization.

#### Immunofluorescence quantification

For staining quantification coverslips were imaged facing down using the EVOS M7000 microscope (Thermo Fisher, Catalog number: AMF7000). To compare control and IONP condition both were imaged using the same imaging protocol, keeping imaging parameters constant. Regions of interest (ROIs) were chosen "blindly" by only considering the DAPI channel, to keep the cell number comparable and avoid introducing experimenter bias. The stainings were quantified depending on the localization of the antibody (cytosol or nucleus) and the experimental model (cell culture). For stainings targeting the whole cell we used a custom CellProfiler$^{TM}$ 4.2.8 pipeline. In brief, DAPI was used as a seeding point (primary object) for cell segmentation and in each staining panel the most cell encompassing staining was chosen to segment the whole cell body (secondary object). For antibodies targeting nuclear proteins (for ex. SOX2) the number of colocalizing positive stainings and DAPI were evaluated. In slice culture stainings fluorescent GBM cells were segmented using a threshold-based ImageJ pipeline to create ROIs. Then pixel intensity of these ROIs was measured in the respective channels. QC plots were produced for each image and evaluated for correct segmentation of nucleus and cells. Then pixel intensity parameters were extracted from each cell body ROI and median pixel intensity was compared between conditions. Statistical evaluation was done using Wilcoxon rank sum test with the addition of effect size testing (Cohen's D).

#### Human organotypic slice culture

Human cortical tissue directly from the operation room was prepared for long term culturing as previously described [87,88]. In short, human cortex (access tissue) removed to access deeper pathologies, was collected immediately after resection from the operation room, submerged in "Preparation Medium" (PM) (Gibco HibernateTM media with 1 mM Gibco GlutaMaxTM, 13 mM Glucose, 30 mM NMDG and 1% Anti-Anti) on ice (0°C ± 4°C) and supplemented with carbogen (95% O2 and 5% CO2). The resected tissue was trimmed and portioned to generate tissue blocks with dimensions of 2cm$^3$. The tissue blocks prepared for sectioning was ensured to contain all the layers of the grey matter as well as a portion of white matter to ensure posthoc orientation of the tissue sections. Tissue blocks are placed in a vibratome chamber (VT1200, Leica Germany) submerged in carbogenated and cooled PM and sectioned to a thickness of 300 µm. Post sectioning, the sections were transferred from the sectioning chamber to the cell culture inserts (Cell culture inserts 0.4 µm & 30 mm, Millicell®, Ref. PICM0RG50) in a 6-well plate, using a fire polished wide-mouthed glass pipette. The tissue sections were cultured in "Growth Medium" (GM) (Neurobasal (L- Glutamine) supplemented with 2% serum free B-27, 2% Antibiotic-Antimycotic, 13mM D+ Glucose, 1 mM MgSO4, 15mM HEPES (Sigma, H0887) and 2mM Glutamax). The sections are incubated at 37°C and 5% CO2 for 7 days.

#### Slice culture injections

One day after sectioning the slices are ready for microsurgical injections. To evaluate the effect if IONPs on tumor progression 12 slices were taken for each patient in the study and divided into control, GBM-injected, IONP-injected and GBM-IONP-injected. A cell suspension with a concentration of 10'000 cells/µL was prepared for injections. 10´000 Cells were inoculated into cortical sections with a 10 µL Hamilton syringe (Hamilton Company, Ref. #80300) into the interface of white and grey matter. For the GBM-IONP-injected condition the injection solution was additionally supplemented with IONPs at 2.5 mg Fe/mL. For the IONP-injected condition GM was supplemented with IONPs at a concentration of 2.5 mg Fe/mL and injected as above. The slices received a medium change every 24h and are incubated for 6 days after injection. Tumor growth was recorded by means of fluorescence imaging (EVOS M7000 microscope) using the same imaging protocol to allow for comparative tumor progression analysis between conditions over time.

#### Slice culture tumor progression analysis

Image analysis is done using FIJI. To track tumor proliferation a combination of area and intensity of fluorescent GBM cells was monitored over the culturing period. Area alone (showing the extent of tumor growth in the x and y plane) is not sufficient to accurately reflect tumor progression as it isn't able to capture growth in the z plane. Tumor growth in the z plane however is reflected in an increase in fluorescent intensity due to overlapping cell´s fluorescence. Every day of culturing (including day of injection) the slices were imaged under brightfield (to create masks of the slice) and under fluorescence (for tumor cell segmentation). The fluorescent images were cropped using the masks to only include cells that grow within the slice (cells growing into the slice from the edges are excluded). Background subtraction with a rolling ball radius of 90 pixels is performed and the image is thresholded to retain only the tumor cells. Finally, the “Analyze Particles” is run with a particle size of 0-infinity to select all thresholded cells (size filtering will be done at a later stage in the analysis) (**Suppl. Fig. S4A**). The detected areas are used as regions of interest (ROIs) and used to measure area and intensity in the original raw image. These two metrics were multiplied to produce the tumor progression score as an approximation of the tumor´s growth in 3 dimensions. All values are normalized to the tumor progression score of the day of injection.

#### Slice culture tumor cell and cluster

To have a more nuanced analysis of tumor growth and cluster formation in the Human organotypic slice culture, detected objects from the **Slice Culture Proliferation analysis** were classified based on their size. To separate the data into debris, individual cells, small/medium/large clusters we evaluated the size of individual cells. For this we manually selected 150 individual cells from day 2 (50 cells from one slice of each patient) and plotted the distribution (**Suppl. Fig. S4B**). Cells were represented by objects with areas between 135 and 2224. This way we defined anything smaller to be cellular debris (excluded from further analysis), anything within this range to be cells and we arbitrarily chose to divide the larger objects into small (2224-5000 $px^2$), medium (5000-10000 $px^2$) and large (10000- $\infty$ $px^2$) clusters (**Suppl. Fig. S4C**). We then followed the changes of object numbers over time. All object counts are normalized to day 0 (day of injection).

#### Slice culture invasion

To quantify tumor invasion specifically beyond tumor progression, we assessed the extent to which cancer cells infiltrate healthy, non-tumorous tissue. We use the location of tumor cells extracted in **Slice Culture Proliferation analysis** to estimate the extent of tissue invasion. We generated a concave hull around all detected cells using the R package “concaveman”. This hull represents the minimum area encompassing the spatial distribution of tumor cells, providing an approximation for the area of invasion over time. All values are normalized to the area of invasion at the day of injection.

#### Slice culture tissue colonization

As an additional measure of the tumors potential to grow and thrive in the cortical tissue we examined tissue colonization. This combines extracted metrics from the **Slice culture proliferation analysis** and the **Slice culture invasion analysis**. The resulting metric represents the tumor´s ability to proliferate in and colonize the area it has invaded. For this we calculate for any given day the fraction of invaded area (from **Slice culture invasion analysis**) that is occupied by the tumor (from **Slice culture proliferation analysis**) over the course of the experiment.

### Supplementary figures

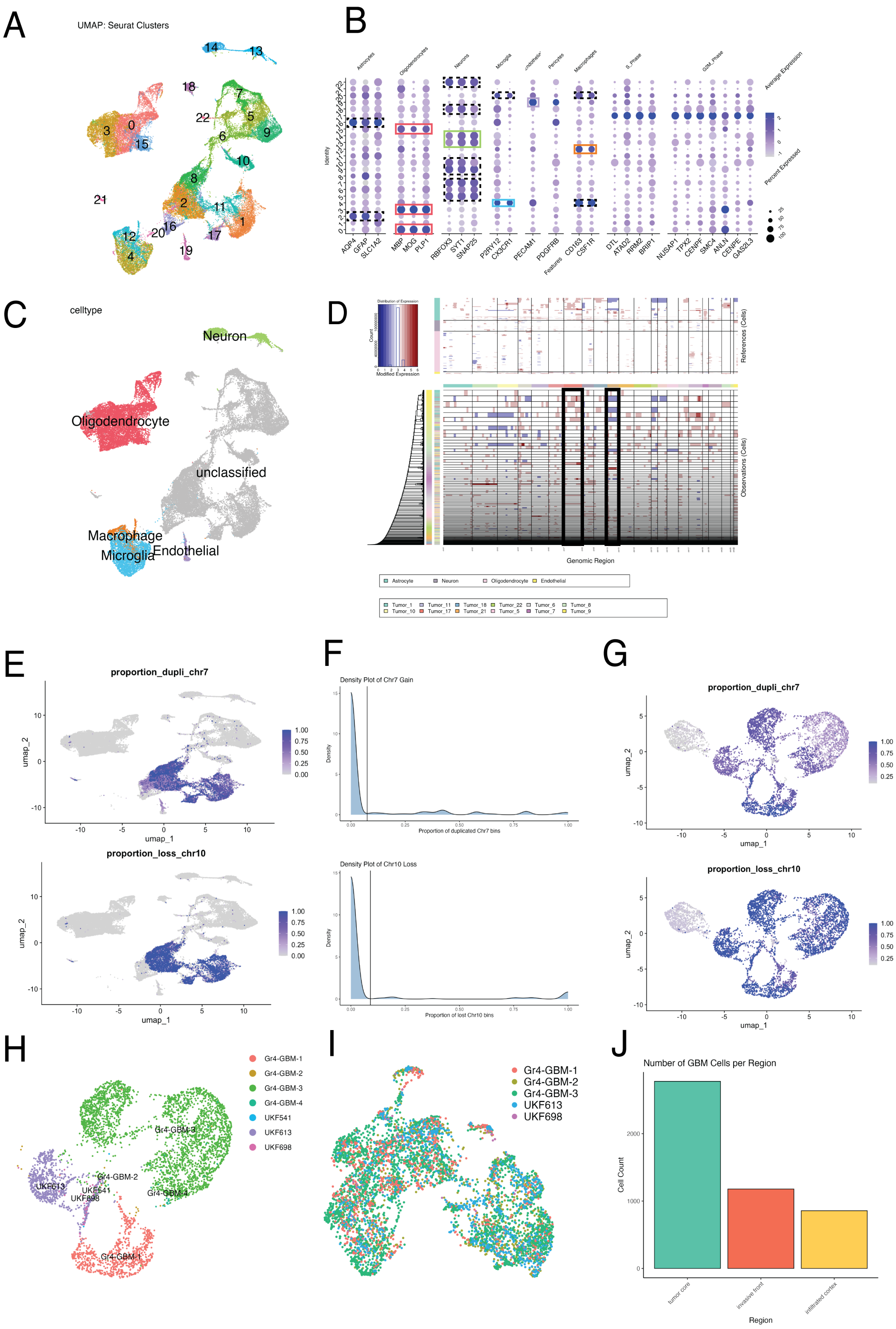


Figure S1: Single nucleus RNA sequencing cell annotation. (A) UMAP showing the seurat clusters of the whole integrated data set. (B) Dot plot showing the expression of canonical cell marker genes and cell cycle genes in the different seurat clusters. Boxes show the clusters suspected cell type, dashed boxes indicate clusters that do show canonical gene expression but were conservatively left unlabbeled for later CNV analysis. (C) Conservative cell annotation leaving all cells that cluster together around the suspected GBM cells as “unclassified”. (D) iCNV plot after running the analysis on the “unclassified” cells. Black boxes indicate the chromosomes of interest, GBM is characterized by gains in chromosome 7 and losse in chromosome 10. (E) UMAP with overlaid scores related to the classic GBM CNVs. (F) Thresholds chosen for the CNVs based on which GBM cells were defined. (G) After subsetting the GBM cells based on the CNV thresholds in (F), cells were again plotted in a UMAP and colored based on their CNV profiles. We can clearly see that one cluster stands out with combined low scores for gains in chromosome 7 and losse in chromosome 10. This cluster was excluded from the GBM data set. (H) UMAP of GBM cells before integration and (I) after integration colored based on patient (note two patients were removed due to low cell numbers for accurate integration). (J) Numbers of GBM cells based on region.

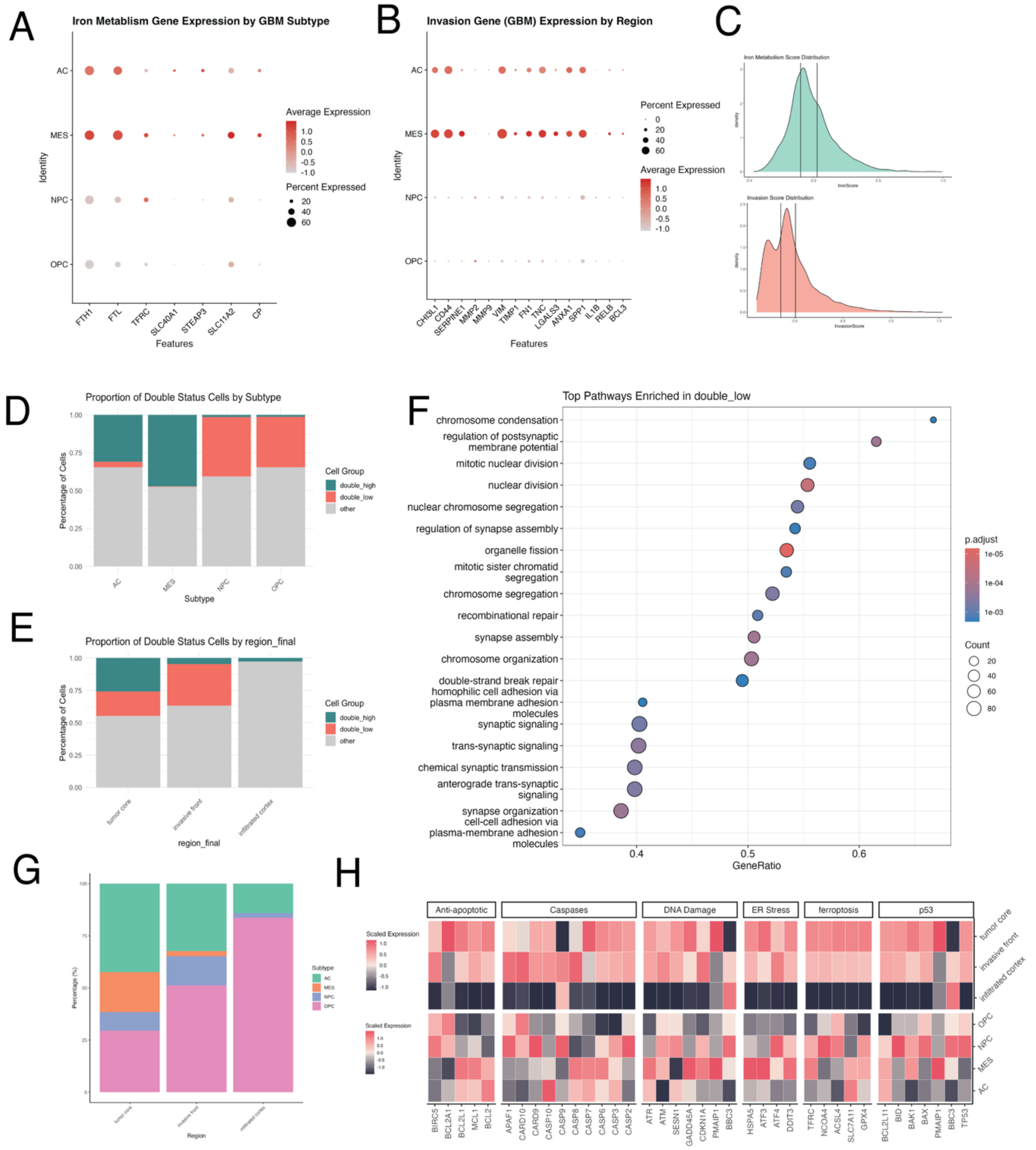


Figure S2: Characterization or cell´s iron and invasion related gene expression. (A) Dot plot showing the expression of iron related genes depending on the GBM subtype, as defined by Neftel et al. 2019. (B) Dot plot showing the expression of invasion related genes depending on the GBM subtype. (C) Tertiles chosen across the expression spectrum of iron metabolism (top) and invasion related genes (bottom) to define cells into low, medium and high. (D & E) Bar plots showing the proportion of double_high and double_low cells depending on the GBM subtype and region respectively. (F) Pathway analysis showing the gene sets that are enriched in the double_low cells. (G) Bar plot showing the proportion of GBM subtypes among malignant cells in the different sampling regions. (H) Heatmap showing the expression of genes related to apoptosis in the different regions and GBM subtypes.

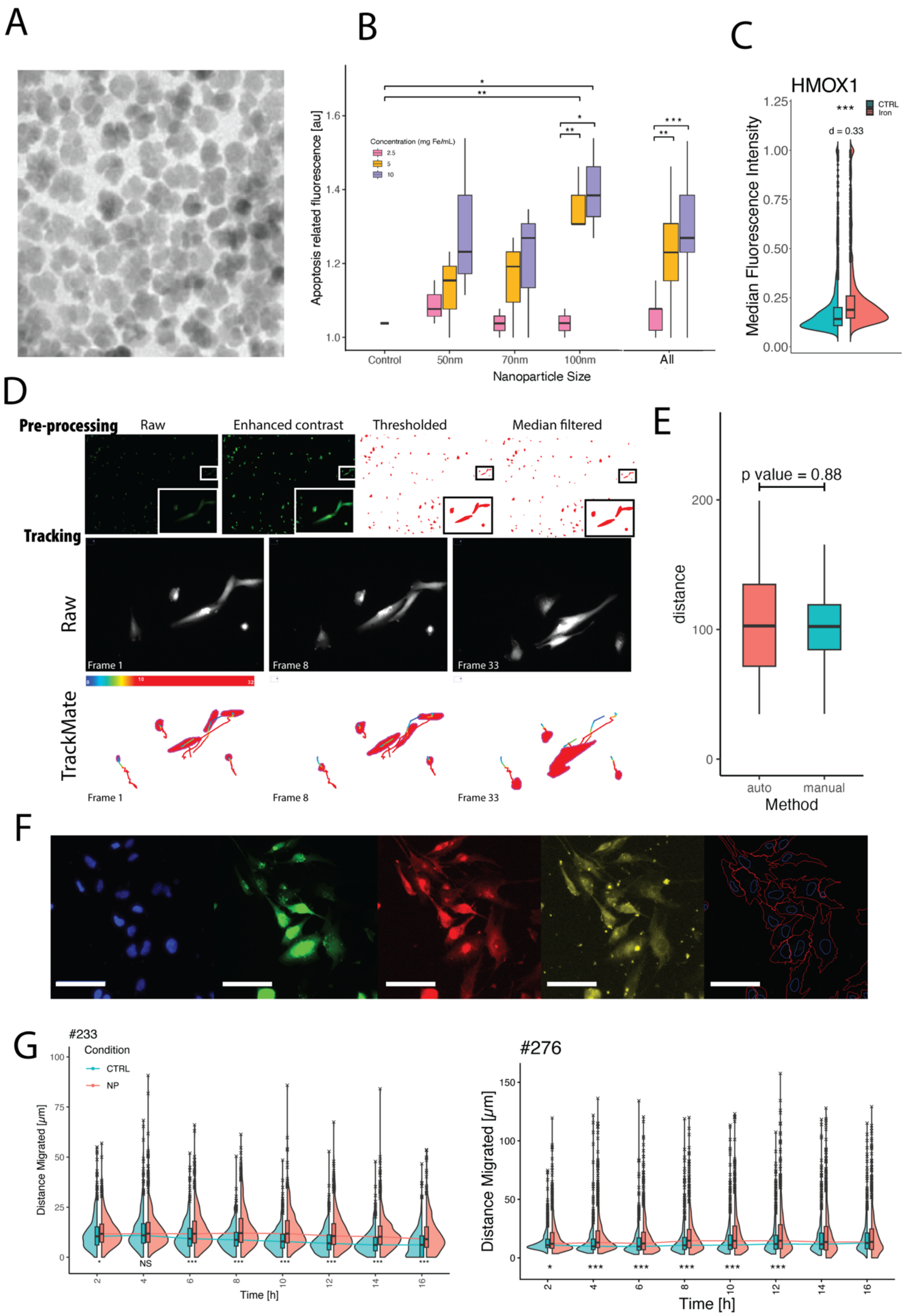

A
B
Concentration (mg Fe/mL)
2.5
5
10
Apoptosis related fluorescence [au]
Control
50nm
70nm
100nm
All
Nanoparticle Size
C
HMOX1
CTRL
Iron
d = 0.33
Median Fluorescence Intensity
D
Pre-processing
Raw
Enhanced contrast
Thresholded
Median filtered
Tracking
Raw
Frame 1
Frame 8
Frame 33
TrackMate
Frame 1
Frame 8
Frame 33
E
p value = 0.88
distance
auto
manual
Method
F
G
#233
Condition
CTRL
NP
Distance Migrated [μm]
Time [h]
#276
Distance Migrated [μm]
Time [h]

**Figure S3:** Cell culture iron exposure experiments. (A) Electron microscope image of Synomag®-D 70nm (Source: Core Front Co., Ltd.). (B) TUNEL (Terminal deoxynucleotidyl transferase-mediated dUTP-biotin nick end labelling) assay of GBM cells exposed to different sizes (50, 70, 100nm) iron oxide nanoparticles at different concentrations over 16h, normalized to control. (C) Immunofluorescence stainings comparing HMOX expression in cells that were exposed to IONPs over 24h (red) vs control (blue). Effect size and p-value indicated above the plot. (D) Cell culture tracking using TrackMate. (Top) Pre-processing steps include contrast enhancement, thresholding and median filtering. Shown is the whole image with a zoomed in section (white/black rectangle) for closer inspection. (Bottom) Tracking process allowing for gap closing and splitting under certain conditions. Shown are example raw and matching tracking images from TrackMate. Tracking lines are colored based on frames. (E) Validating the automated tracking by comparing it with manually tracked cells (cells tracked: $n_{auto}$ = 63, $n_{manual}$ = 66). (F) Showing an example of the cell body segmentation used for fluorescence quantification. Example showing DAPI, ZsGreen, CD71 and FTL stainings and the resulting primary (nuclei) and secondary (cell body) segmentation. Scale bar is 100µm. (G) Violin plots showing the differences in the kinetics of the migration response between both cell lines.

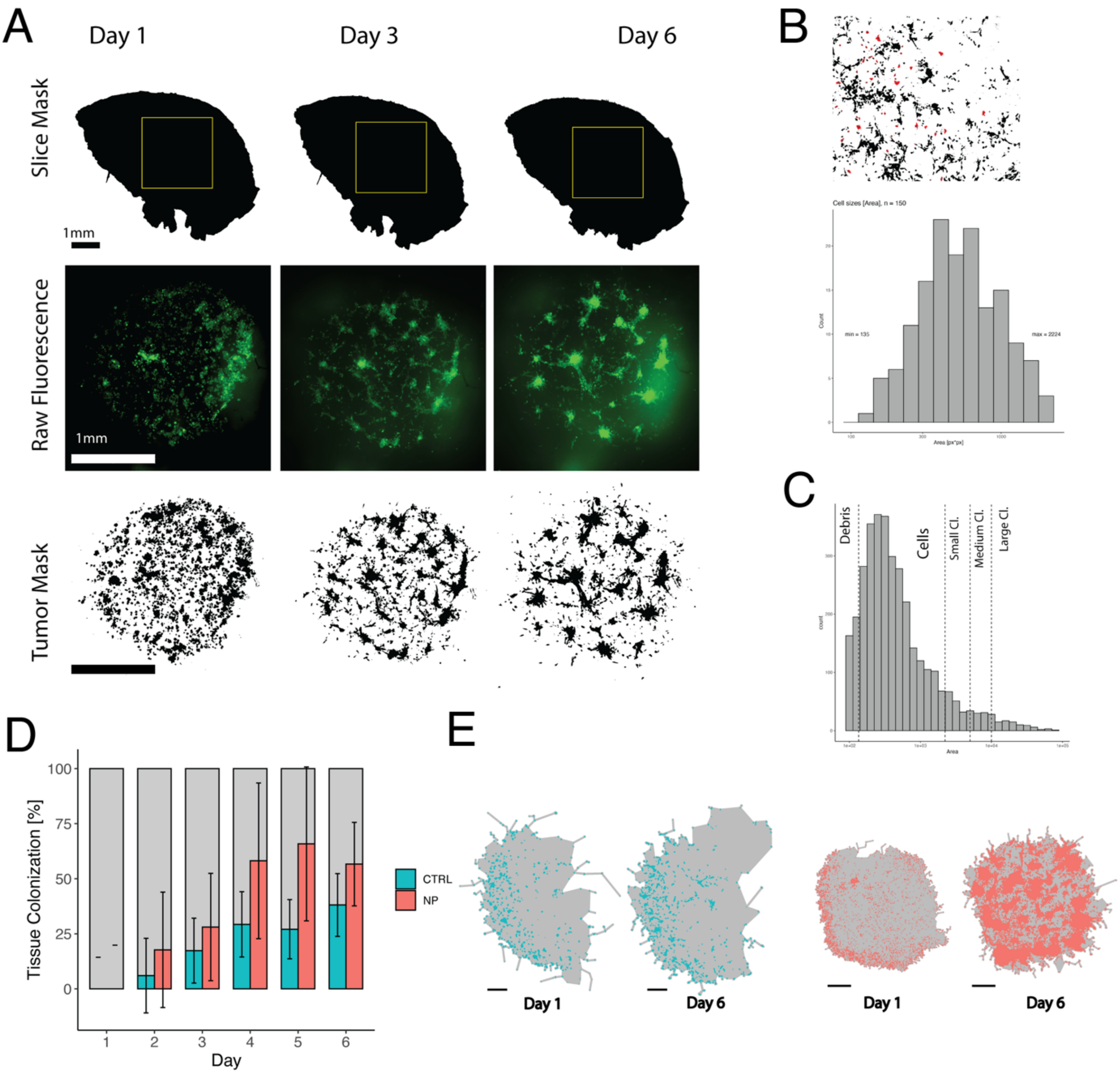


**Supplementary Figure S4: Slice culture.** (**A**) Example of tumor segmentation in the human organotypic slice culture. From top to bottom: Mask from the bright field image of the slice (yellow square shows region shown below), fluorescent image of the tumor growing within the slice and the segmented tumor cells. Segmented cells will be used as ROIs to extract area and intensity from the original fluorescent image. Scale bars are 1mm (**B**) (Top) 50 cells (red) were manually chosen from day 2 of three slices (one from each patient, 150 cells in total) to determine the sizes of cells. (Bottom) Histogram showing the distribution of cell sizes. The minimum size for a cell was determined to be 135 px$^2$ and the maximum 2224 px$^2$. This interval was chosen to define cells. (**C**) Histogram of all objects (n = 24216) detected across the whole experiment. Vertical lines showing the cutoff for defining objects as debris, cells, small/medium/large clusters. (**D**) Bar plot (with error bars) showing tissue colonization over the days, which illustrates how much of the invaded area is colonized by the tumor. Tissue colonization is calculated by dividing the tumor volume (colored by condition) by the area of invasion (grey). It is a metric showing the capability of cells to occupy the area they invaded. (**E**) Two examples showing the tumor colonization from both conditions from day 1 and 6. Scale bar is 500 μm.
